# Machine learning enabled phenotyping for GWAS and TWAS of WUE traits in 869 field-grown sorghum accessions

**DOI:** 10.1101/2020.11.02.365213

**Authors:** John N. Ferguson, Samuel B. Fernandes, Brandon Monier, Nathan D. Miller, Dylan Allan, Anna Dmitrieva, Peter Schmuker, Roberto Lozano, Ravi Valluru, Edward S. Buckler, Michael A. Gore, Patrick J. Brown, Edgar P. Spalding, Andrew D.B. Leakey

## Abstract

Sorghum is a model C4 crop made experimentally tractable by extensive genomic and genetic resources. Biomass sorghum is also studied as a feedstock for biofuel and forage. Mechanistic modelling suggests that reducing stomatal conductance (*g_s_*) could improve sorghum intrinsic water use efficiency (*iWUE*) and biomass production. Phenotyping for discovery of genotype to phenotype associations remain bottlenecks in efforts to understand the mechanistic basis for natural variation in *g_s_* and *iWUE*. This study addressed multiple methodological limitations. Optical tomography and a novel machine learning tool were combined to measure stomatal density (SD). This was combined with rapid measurements of leaf photosynthetic gas exchange and specific leaf area (SLA). These traits were then the subject of genome-wide association study (GWAS) and transcriptome-wide association study (TWAS) across 869 field-grown biomass sorghum accessions. SD was correlated with plant height and biomass production. Plasticity in SD and SLA were interrelated with each other, and productivity, across wet versus dry growing seasons. Moderate-to-high heritability of traits studied across the large mapping population supported identification of associations between DNA sequence variation, or RNA transcript abundance, and trait variation. 394 unique genes underpinning variation in WUE-related traits are described with higher confidence because they were identified in multiple independent tests. This list was enriched in genes whose orthologs in Arabidopsis have functions related to stomatal or leaf development and leaf gas exchange. These advances in methodology and knowledge will aid efforts to improve the WUE of C4 crops.

## INTRODUCTION

Global climatic change is subjecting agricultural regions to elevated atmospheric vapor pressure deficits (VPD; Yuan et al., 2019) and reduced precipitation events (Sheffield and Wood, 2008), thereby giving rise to situations where more water is needed, but less is available. Water use efficiency (WUE; the ratio of carbon gain to water loss) is a key target trait for crop improvement to improve production and sustainable water use (Bailey-Serres et al., 2019; Leakey et al., 2019). C_4_ crops including maize, sorghum, sugarcane, millet and Miscanthus are heavily studied as sources of food, feed, fuel and fiber. However, less research has been directed towards understanding and improving WUE and its component traits in C_4_ crops, possibly because they already achieve high WUE as a result of the CO_2_ concentrating mechanism they possess (DeLucia et al., 2019). Nevertheless, mechanistic modeling suggests that enhancing intrinsic WUE (*iWUE*) by reducing stomatal conductance (*g_s_*) while maintaining rates of net CO_2_ assimilation (*A_N_*) can increase biomass production in C_4_ as well as C_3_ species across a broad range of environmental conditions (Truong et al., 2017; Leakey et al., 2019). These benefits will become greater as atmospheric [CO_2_] continues to rise. Compiling surveys of natural variation in C_4_ species, including grain sorghum (Kapanigowda et al., 2013), demonstrated that *g_s_* could explain substantially more variation in *iWUE* than *A_N_* (Leakey et al., 2019).

Sorghum is a model C_4_ crop made experimentally tractable by extensive genomic and genetic resources (Paterson et al., 2009; Morris et al., 2013). And, biomass sorghum has considerable potential as a biofuel feedstock in addition to being grown for forage (Castro et al., 2015). This study aimed to address key knowledge gaps regarding natural variation in *iWUE* and related traits across diverse biomass sorghum accessions, including evaluation of heritability, environmental effects, trait correlations, and associations between DNA sequence variation or RNA transcript abundance and trait values. *iWUE* was studied alongside its component traits (*A_N_* and *g_s_*) plus stomatal density (SD) and specific leaf area (SLA), because these anatomical and allometric traits are known to influence leaf physiology.

Stomata open and close to regulate the rate of CO_2_ and water vapor exchange between leaves and the atmosphere (Cowan and Farquhar, 1977). These fluxes are also influenced by the size and density of stomata (Franks and Beerling, 2009; Dow et al., 2014). Empirical data shows that SD is positively correlated with *g_s_* in a number of species (Anderson and Briske, 1990; Pearce et al., 2006). Molecular mechanisms controlling stomatal morphology and patterning have been elucidated in *Arabidopsis thaliana* (Chater et al., 2017). This has been combined with understanding of how *g_s_* and WUE are linked to stomatal physiology to develop C_3_ plants with improved WUE. For example, the expression of species-specific orthologs of the *A. thaliana EPIDERMAL PATTERNING FACTOR 1* (*EPF1*) gene has been targeted to reduce *g_s_* through reduced SD, thereby improving WUE in barley (Hughes et al., 2017), rice (Caine et al., 2019; Mohammed et al., 2019), wheat (Dunn et al., 2019) and poplar (Wang et al., 2016). The majority of cultivated crops are grasses (Leff et al., 2004). Stomatal morphology and development in grasses is markedly different from that of dicotyledonous species, e.g. *A. thaliana*, and reflects specific selective pressures (Hetherington and Ian Woodward, 2003). Consequently, whilst in some instances the molecular underpinnings of these traits are conserved between *A. thaliana* and grasses (Hughes et al., 2017; Caine et al., 2019; Dunn et al., 2019; Mohammed et al., 2019), emerging evidence suggests the biological functioning of key stomatal genes can be divergent between the lineages (Raissig et al., 2017; Abrash et al., 2018). As such, improving our understanding of grass-specific genes that regulate stomatal development and patterning will expedite efforts to improve WUE in crops. The need to address this knowledge gap is greatest in C_4_ species.

SLA is the ratio of leaf area to leaf mass, which combines information on leaf thickness and leaf density (John et al., 2017). It is a key trait in the leaf economic spectrum than influences many traits including photosynthesis, respiration, leaf construction costs, leaf life span, canopy light interception and growth rates (Wright et al., 2004). Despite its importance, and that it can be measured easily, efforts to understand the genetic architecture of the trait through quantitative trait loci mapping or GWAS have been limited (Yin et al., 1999; El-Lithy et al., 2004; Trachsel et al., 2010). But, correlations between SLA and SD have been observed in response to varying water supply (Xu and Zhou 2004) and across intraspecific variation associated with adaptation to aridity (Carlson et al., 2016). The genetic and environmental control of SLA and its relationship to SD in C_4_ species is especially poorly understood.

Efforts to discover the genetic basis of traits that influence the sustainability and resilience of crop productivity, including *iWUE*, are constrained by bottlenecks in both phenotyping as well as discovery of associations between trait variation and DNA sequence variation or gene expression (Yang et al., 2020). Automation, remote sensing and machine learning are increasingly being used to accelerate the measurement and/or quantification of key ecophysiological traits (e.g. Atkinson *et al.* 2017; Banan *et al.* 2018; Feldman *et al.* 2018; Qiao *et al.* 2019). Optical tomography has been proposed as a method for imaging cell patterning on leaf surfaces that is much more rapid than traditional methods of epidermal peels or imprinting (Haus et al., 2015). Identifying and counting stomatal complexes on the epidermis is the most time-consuming aspect of screening SD. A number of machine learning tools have been proposed for counting stomata (e.g. Fetter et al., 2019; Li et al., 2019; Sakoda et al., 2019). However, proof of concept is still required for the use of optical tomography and an automatic stomatal counting tool suitable for use across the phenotypic variation associated with diverse genotypes of a grass species.

Genome-wide or transcriptome-wide association studies (GWAS and TWAS) are popular methods that can identify genomic regions or genes for which variation in DNA sequence or gene expression is associated with quantitative variation in a trait of interest (Tian et al., 2011; Hirsch et al., 2014; Xu et al., 2017). The challenges associated with phenotyping traits associated with *iWUE* in C_4_ crops mean that these methods have only been applied in a limited number of cases (Ortiz et al., 2017; Feldman et al., 2018, Ellsworth et al., 2020). But, even when phenotypic data is readily available, association studies are often challenging because many traits are highly polygenic, where a large number of genes each exert a weak effect on the trait (Zhu et al., 2008). Larger mapping population sizes can improve statistical power to counteract this problem. But, multiple testing at many single nucleotide polymorphisms across the genome also creates a significant risk of false positive results. Validating the function of candidate genes via reverse genetics remains the gold standard, but is extremely slow. Approaches that can increase confidence and efficiency of identification of candidate genes from association studies are therefore important. One simple approach is to prioritize genes identified in multiple independent tests. Alternatively, GWAS can be supplemented by TWAS. Most recently, proof-of-concept for applying Fisher’s combined test to integrate GWAS and TWAS was provided by demonstrating how it increased the efficiency with which known causal genes could be “re-discovered” for well-studied maize kernel traits (Kremling et al 2019). However, the application of the method to address knowledge gaps for traits such as *iWUE* is untested.

In summary, to address knowledge gaps about the physiology and genetics of natural variation in *iWUE* in C_4_ grasses, this study evaluated a diverse population of 869 biomass sorghum accessions grown in replicated trials over two growing seasons. To achieve this goal, a set of novel tools were developed, tested and integrated. To break the phenotyping bottleneck for SD, optical tomography was adapted and tested as an imaging technology and a custom machine learning software platform was developed to automatically identify and count stomatal complexes. This was combined with a rapid method to measure leaf-level gas exchange and specific leaf area (SLA). Trait correlations were evaluated and genes putatively underlying genetic variation in *iWUE* and related traits were identified through GWAS, TWAS and an ensemble association mapping approach.

## RESULTS

### Growing season climate

A diversity panel of 869 photoperiod-sensitive sorghum accessions (Figure S1; Table S1) was grown at field locations within a five-km radius in 2016 (N=2; Fisher and Energy Farms) and 2017 (N=2; Maxwell and Energy Farms). Mean daytime maximum temperature was similar between 2016 (28.9 °C) and 2017 (28.8 °C). But, compared to the average growing season rainfall of 396 mm (Gelaro et al., 2017), 2017 was dry (174 mm) and 2016 was wet (529 mm; Figure S2).

### High-throughput phenotyping metrics

A high-throughput approach for measurement of photosynthetic gas exchange (*g_s_*, *A_N_*, *iWUE*, and the ratio of intracellular to atmospheric CO_2_ concentration (*c_i_*/*c_a_*)) along with tissue sampling for SLA and SD was performed on ~220 leaves per day, allowing two leaves per replicate plot of every genotype in the population to be sampled through 9-10 days of work for each replicate field in a given year.

Optical topometry (OT) was used to rapidly image 4-6 fields of view (FOV) from the abaxial surface of 4169 leaves in 2016 and 3211 leaves in 2017 without the need for sample preparation beyond adhesion to microscope slides with double-sided tape (~250 FOVs per day per OT microscope; Figure 1A). High-throughput computing resource allowed SD to be assessed for each of the 33,355 FOVs in <24 hours using a convolutional neural network that was trained to identify stomatal complexes in a rotationally invariant manner (Figure 1B; Figure S3). In contrast, based on recent experience, manual counting of this image set would take an estimated 80 person-days. The median SD per leaf generated by this machine-vision platform was significantly positively correlated (R^2^ = 0.72, p<0.001) with the median SD per leaf from human counting of 228 randomly selected ground truth samples (Figure 1C). Although, there was a bias towards overestimation of SD by the computer as a result of a low rate of false positive identification of cells as stomatal complexes, especially on leaves with lower stomatal density.

**Figure 1.**
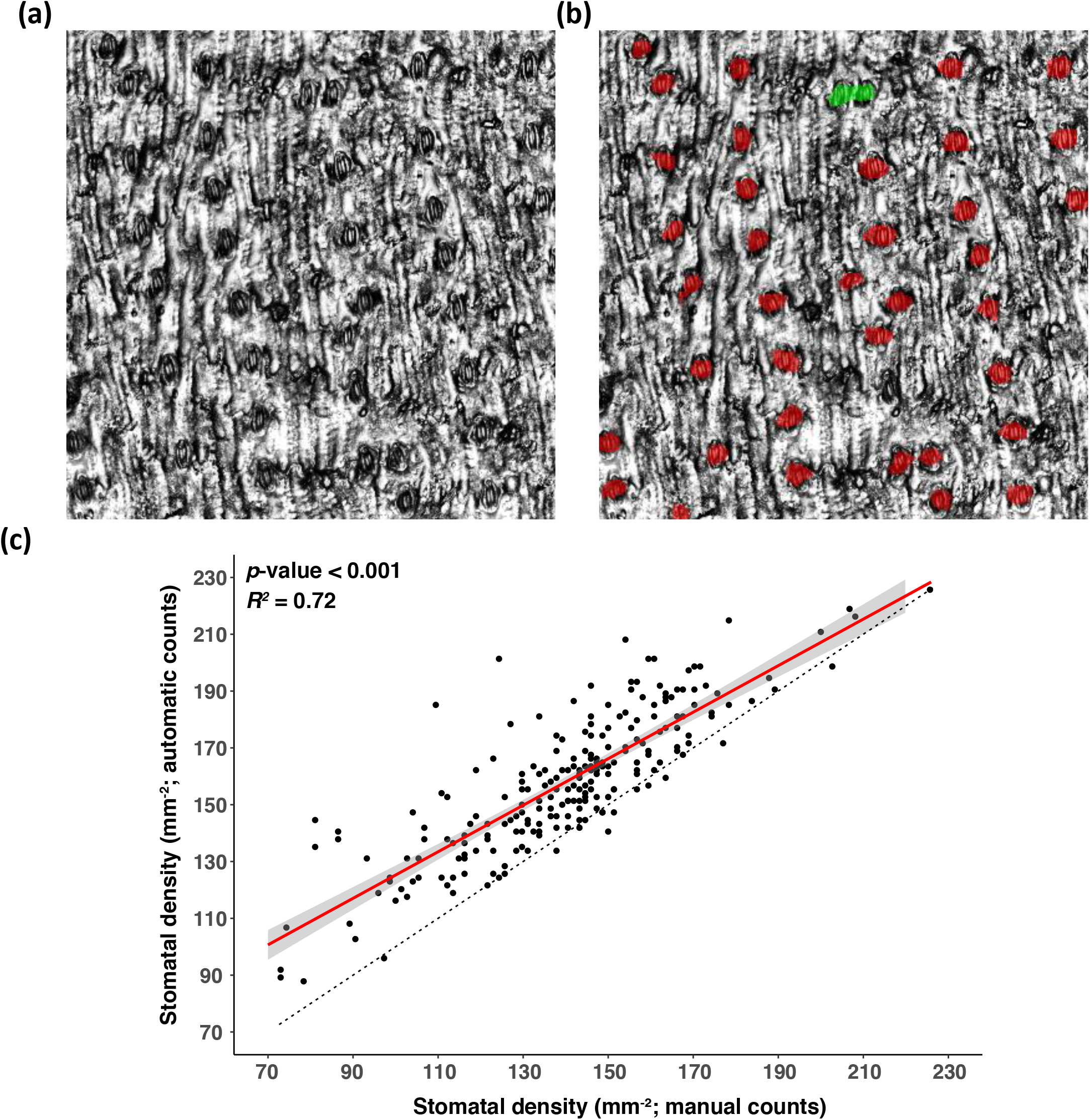
Demonstration of stomatal counting algorithm. (a) Reflective intensity layer of an optical topometry (OT) measurement of the abaxial epidermis of a sorghum leaf section. (b) OT measurement overlaid with automatic detection of stomata (red). Automatically detected stomata in close proximity are highlighted in green. (c) Association between median stomatal density of samples where stomates have been both manually and automatically counted. 4-6 fields of view were used to calculate the median values for each genotype by manual and automatic methods. A linear model regressing automatic counts on manual counts is fitted (red) and the standard error of the model is shown (gray). The associated *p-value* significance threshold and *r^2^* value of the model are inset.

### Natural variation of WUE associated traits

Under the drought conditions of 2017, SD was 40% lower on average than in wetter conditions of 2016 (Figure 2A). The range of trait variation was also 26% less in 2017 than 2016. In contrast, SLA was 24% greater, on average, in 2017 than 2016. Again, this was associated with less variation within the population in the drought year (Figure 2B). Leaf photosynthetic gas exchange was only measured in 2017. On a relative basis, the observed variation in trait values was greatest for *g_s_* (0.17 to 0.41 mol m^−2^ s^−1^) and *A_N_* (19.8 to 35.0 μmol m^−2^ s^−1^), moderate for SD and SLA in the same year, and least for *iWUE* (108 - 151 μmol mol^−1^) and *c_i_*/*c_a_* (0.40 – 0.52; Figure 2C-F). Plant height was strongly correlated between 2016 and 2017 (r = 0.94; Figure S15). Therefore, adjusted genotypic means were calculated for plant height from a mixed model incorporating data from both 2016 and 2017 at the growth stage where heritability was highest (height-joint; Figure 2G). The mean height-joint was 302 cm and varied within the population from 143 to 381 cm. Plant biomass was only measured in 2017. Mean biomass production was 28.7 t ha^−1^ (Figure 2H).

**Figure 2.**
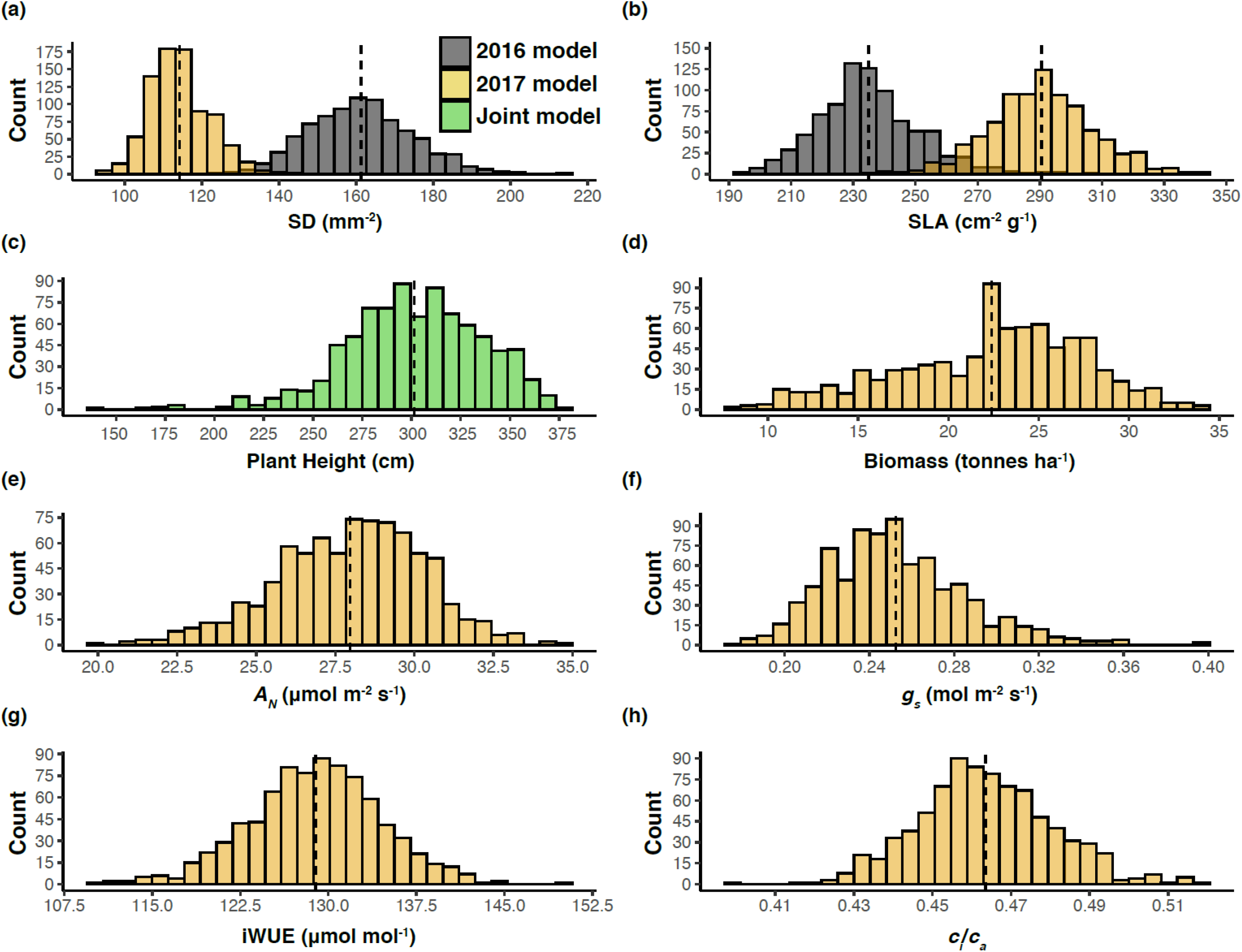
Histograms of variation in adjusted means of (a) Stomatal density (SD), (b) specific leaf area (SLA), (c) plant height (*g_s_*), (d) above-ground biomass (*A_N_*), (e) net photosynthesis (*A_N_*), (f) stomatal conductance (*g_s_*). (g) intrinsic water use efficiency (*iWUE*), and (h) ratio of intracellular to atmospheric CO2 concentration (*c_i_*/*c_a_*). The dashed vertical lines denote the population mean.

SD in 2016 was significantly correlated with SD in 2017 to a moderate degree (r = 0.36; Figure 3, see Table S2 for details). Variation among the population in SLA was slightly more consistent across wet and dry growing conditions, resulting in a stronger correlation in SLA between 2016 and 2017 (r = 0.46; Figure 3). Plant height was also positively correlated with above-ground biomass production in 2017 (Figure 3).

**Figure 3.**
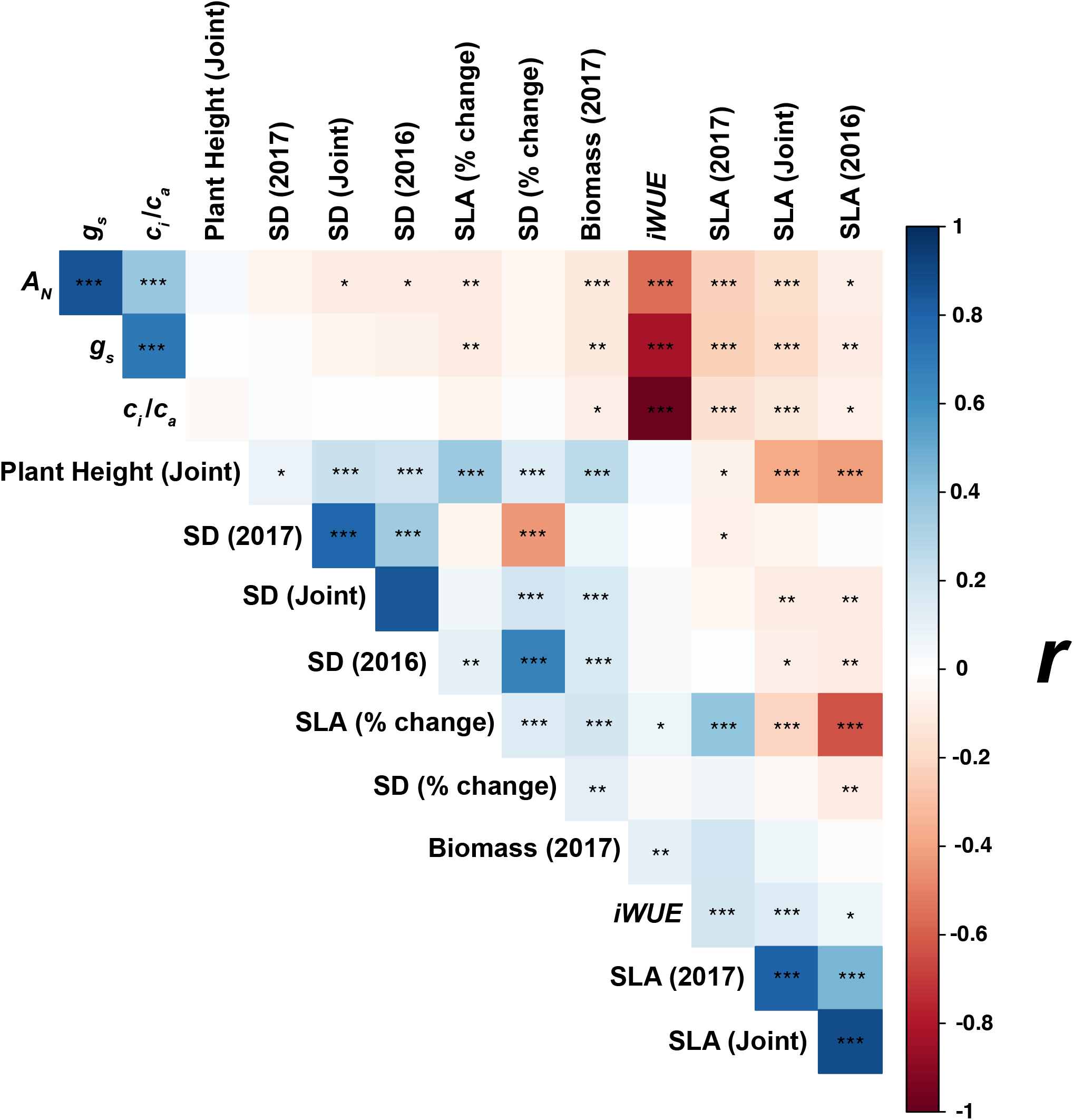
Correlogram for all measured parameters. Where appropriate, the associated model from which predicted means were extracted is indicated in parenthesis. Traits are arranged according to the angular order of eigenvalues. The color of each individual square describes the Pearson’s correlation coefficient (*r*) of each pairwise interaction. Significant correlations are denoted at the level of 0.001 (***), 0.01 (**), and 0.05 (*). Correlations between traits extracted from each individual environment model are listed in Table S2.

SD in 2016, 2017, and when genotypic means for SD were calculated in a joint model incorporating data from both years (SD-joint), were not significantly correlated with *g_s_*, *iWUE* or *c_i_*/*c_a_* (Figure 3). However, SD in 2016 and SD-joint were weakly negatively correlated with *A_N_* in 2017. And, SD was weakly negatively correlated with SLA within each of the two growing seasons. All three SD traits were positively correlated with height. Similarly, SD in 2016 and the SD-joint were positively correlated with biomass production in 2017. The relative change in SD between growing seasons varied from −9 to −47 %, and was weakly, positively correlated with biomass production and height (Figure 3).

*A_N_* and *g_s_* were positively correlated with each other (Figure 4A). *A_N_* and *g_s_* were both negatively correlated with *iWUE*, but *g_s_* explained more than twice as variation in iWUE (*R^2^* = 0.69) than *A_N_* (*R^2^* = 0.32; Figure 4B, C). *iWUE* was weakly positively correlated with biomass production in 2017, but was not correlated with height. *iWUE* was also positively correlated with all SLA traits, with the relationship being strongest for SLA measured in the same growing season as the photosynthetic gas exchange i.e. 2017.

**Figure 4.**
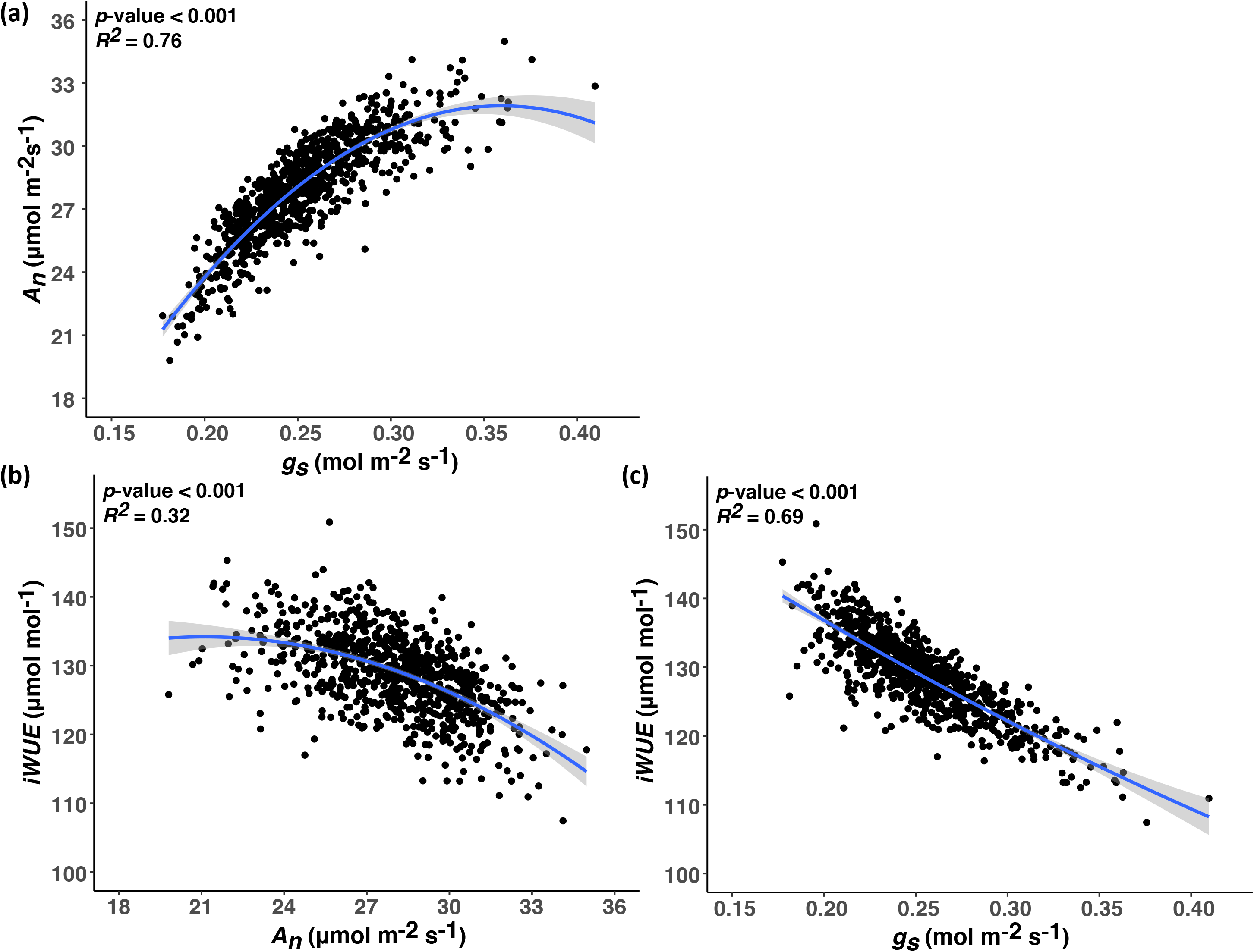
Relationships between gas exchange-related traits. All data are adjusted means. (a) Relationship between stomatal conductance (*g_s_*) and net photosynthesis (*A_n_*). (b) Relationship between An and intrinsic water use efficiency (*iWUE*). (c) Relationship between *g_s_* and *iWUE*. For each relationship, a second-order polynomial model regressing y on x is fitted (blue) and the associated standard error of the model is highlighted (gray). The associated *p-value* significance threshold and *r^2^* values for each model are inset.

To varying degrees in each year, SLA in 2016, 2017 and SLA-joint were all negatively correlated with *A_N_*, *g_s_*, *c_i_*/*c_a_*, SD, and height. The relative change in SLA between growing seasons varied from −2 to +34 %, and was positively correlated with the relative change in SD between growing seasons as well as height, SD in 2016 and iWUE.

### Genetic basis of WUE associated traits

Generalized heritability, was relatively high for SD and SLA (Figure 5). However, the heritability for SD in 2017 (0.50) was lower than in 2016 (0.68) and for SD-joint (0.69). In contrast, heritability for SLA-joint (0.80) was greater than for the individual years of 2016 (0.68) and 2017 (0.71). Leaf-level gas exchange traits demonstrated low to moderate heritability (*g_s_* = 0.44, *A_N_* = 0.42, *iWUE* = 0.31, *c_i_*/*c_a_* = 0.26).

**Figure 5.**
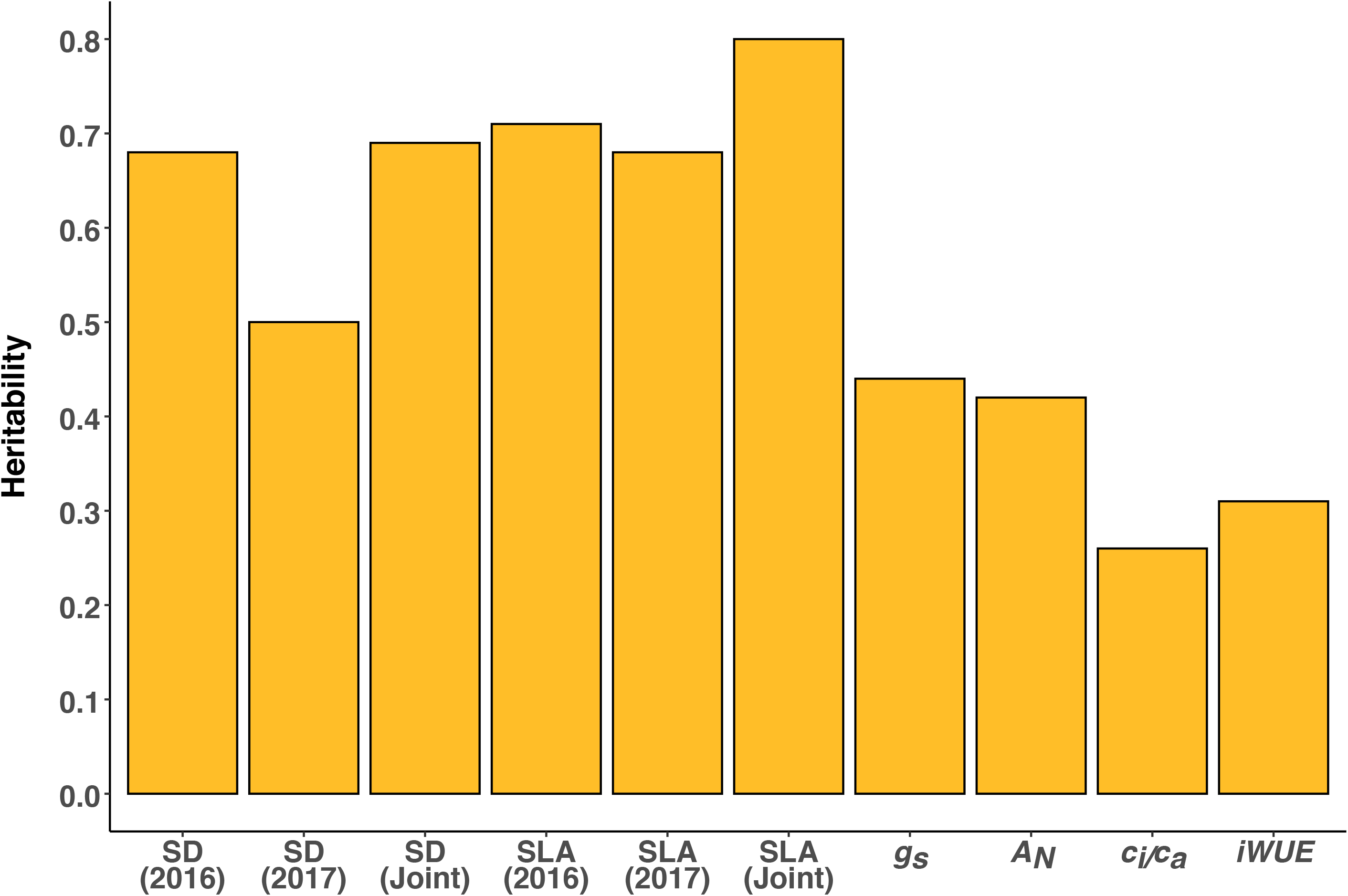
Bar plot of heritabilities of stomatal density (SD), specific leaf area (SLA), stomatal conductance (*g_s_*), net photosynthesis (*A_N_*), the ratio of intracellular to extracellular CO2 (*c_i_*/*c_a_*), and intrinsic water use efficiency (*iWUE*). Models combining individual and joint yearly data were used to estimate heritability for SD and SLA. Gas exchange traits were only measured in 2017.

A three-tiered approach for genetic mapping was used to identify candidate genes underlying the variation observed for the WUE-associated traits under study. Adjusted genotypic means from each linear mixed model for each trait were used for a genome-wide association study (GWAS; e.g. Figure 6A). The genes within linkage disequilibrium (LD) of the most statistically significant 0.1% of GWAS SNPs (Table S3) were then identified (Table S4). The number of independent genes identified per trait varied from 475 for SLA in 2016 to 656 for *A_N_* in 2017 (Figure 6F; Figures S4 – S13). The 1% of gene transcripts that had the most statistically significant associations with a given trait (Table S5; Figure 6B and C; Figures S4 – S13) were identified in transcriptome-wide association studies (TWAS) performed independently for the shoot growing point (GP; 195 genes per trait) as well as the developing third leaf (3L; 167 genes per trait). The p-values for these lists of “top hit” genes identified by GWAS and TWAS were integrated via the Fisher’s combined probability test to identify candidate genes that the two orthogonal tests suggest underlie the observed phenotypic variation across the population (Table S6; Figure 6D and E; Figures S4 – S13).

**Figure 6.**
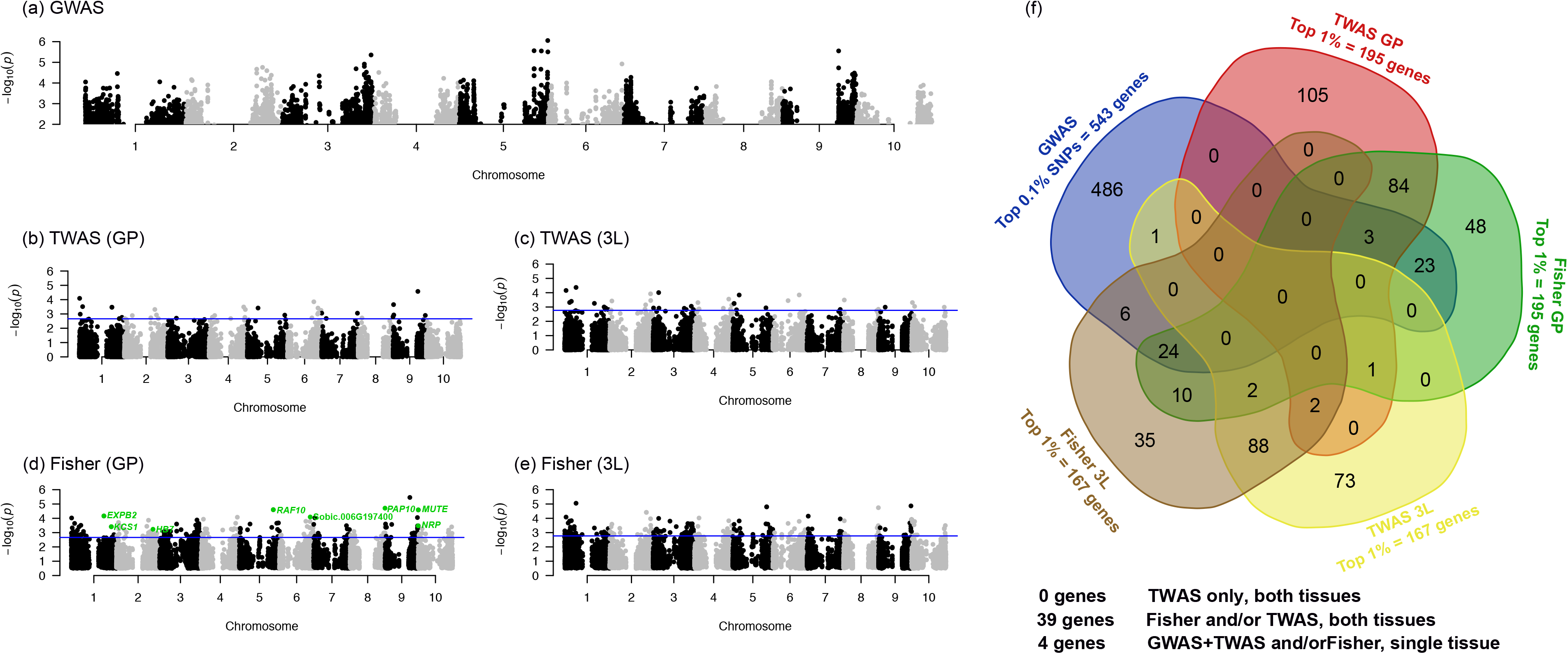
Manhattan plots for stomatal density (SD) in 2016 for: (a) GWAS; (b) TWAS in growing point (GP) tissue; (c) TWAS in the third leaf (3L); (d) Fisher’s combined test results in GP tissue; (e) Fisher’s combined test in 3L tissue; and (f) a five-way Venn diagram highlighting where genes within the top sets of all mapping approaches for SD-2016 are consistently identified. Blue lines indicates the threshold for the genes with the top 1% of −log_10_ (*p-values*). Genes with known or putative roles in stomatal development are highlighted in green. Table S8 for gene lists related to each test, tissue, year and trait.

Candidate genes identified in two or more independent tests are less likely to be false positives i.e. more likely to be associated with genetic variation in the traits of interest. Therefore, the consistency in results was tested: (a) across test types for a single trait (Figure 7A); and (b) across key trait groups, years or test-types (Figure 7A). Between 37 and 59 candidate genes were identified with high confidence for a given trait, based on being identified in at least two independent tests (Figure 7A, Figure 6, Figures S4-S12, Tables S7 and S8). This criterion was most consistently met when the tests integrated data about trait associations with both DNA sequence and RNA transcript abundance, and did so for transcript abundance in both the developing leaf (3L) and growing point (GP) (Figure 7A). For example, 48 genes in total met this criteria for SD in 2016 (Figure 6F) by being consistently identified by: both Fisher’s tests (12 genes), both Fisher’s tests plus both TWAS tests (2 genes), both Fisher’s tests plus one TWAS test (3 + 1 genes), both Fisher’s test plus the GWAS (29 genes), or a Fisher’s test and both TWAS tests (1 gene). In addition, a moderate number of high confidence genes were identified when the tests integrated data about trait associations with DNA sequence and RNA transcript abundance in a single tissue (Figure 7A). For example, 7 genes met this criteria for SD in 2016 by being consistently identified by the GWAS and TWAS (1 gene) or GWAS, TWAS and Fisher’s test (4 + 2 genes) for a given tissue (Figure 6F). The smallest number of “high confidence” genes were found by being identified in TWAS tests for both tissues, without evidence for genotype to phenotype associations from GWAS (Figure 7A). For example, 2 genes met this criteria for SD in 2016 (Figure 6F). These patterns were consistent for all the traits tested. When compiled across all the leaf traits, these multiple independent tests identified 394 unique candidate genes for associations with trait variation for RNA plus DNA, or from both tissues where the transcriptome was tested (Figure 7B, Table S7 and S8).

**Figure 7.**
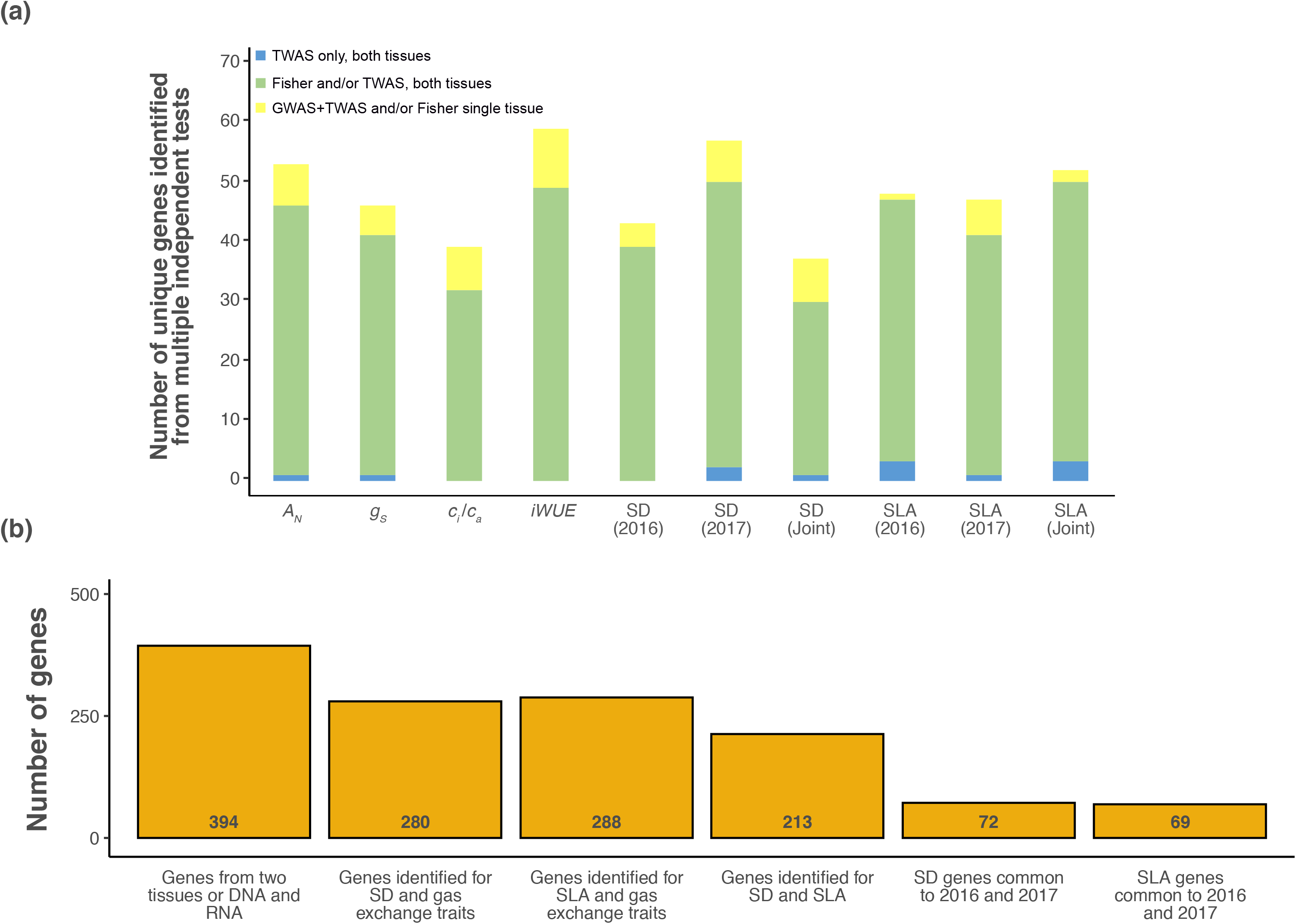
(a) Barplot of the number of unique genes identified with higher confidence as potentially underlying variation on a trait-by-trait basis for net photosynthesis (*A_N_*), stomatal conductance (*g_s_*), the ratio of intracellular to extracellular CO2 (*c_i_*/*c_a_*), and intrinsic water use efficiency (*iWUE*), stomatal density (SD), and specific leaf area (SLA). Higher confidence genes were defined as those identified from multiple tests representing independent evidence from either: TWAS only, but in both tissues (blue fill); Fisher’s combined test and/or TWAS in both tissues (green fill); or GWAS plus TWAS or Fisher’s combined test in one tissue (yellow fill). (b) Barplot of the number of unique genes consistently identified in multiple independent tests across different traits or growing seasons. For reference, the total number of unique genes identified by a parallel trait by trait approach (394, see panel a) is presented in the first bar. See Table S8 for gene lists related to each test, tissue, year and trait.

Candidate genes were also consistently identified by two or more tests that spanned key trait groups (Figure 7B). For example, 213 genes were independently identified by tests for both SD and SLA. 280 genes were independently identified by tests for both SD and photosynthetic gas exchange traits. 288 genes were independently identified by tests for both photosynthetic gas exchange traits and SLA. Comparing across independent tests in separate growing seasons, 72 genes associated with variation in SD were consistently identified in 2016 and 2017. While 69 genes associated with variation in SLA were identified in both years.

In at least 75 cases, the orthologs in Arabidopsis of candidate genes identified here are annotated by TAIR (www.arabidopsis.org) as having some function related to leaf development or WUE (Table S9 and references therein). For example, AT3G06129 (MUTE) shares the greatest sequence similarity with Sobic.009G260200 identified for SD, and enocdes a bHLH protein that controls meristemoid differentiation during stomatal development (Kim et al 2012). AT1G51660 (MAPK4) is most similar to Sobic.004G323600 identified for *g_s_* and is disease resistance protein involved in ABA-regulated stomatal movements (Hettenhausen, Baldwin & Wu 2012; Witoń *et al.* 2016). AT4G00430 (PIP1;4) is most similar to Sobic.006G176700 identified for *A_N_* and is CO2 transporter involved in photosynthetic metabolism (Li et al., 2015).

## DISCUSSION

The trade-off between carbon gain and water use is a fundamental constraint for crop productivity and environmental resilience (Bailey-Serres et al., 2019; DeLucia et al., 2019; Leakey et al., 2019). More specifically, improving WUE is recognized as a means to enhance the utility of sorghum as a biofuel feedstock (Mathur et al., 2017; Meki et al., 2017). Nevertheless, understanding of genetic variation in traits that underlie iWUE in C_4_ grasses is poor even after more than a century of WUE research (Briggs and Shantz 1917; Leakey et al., 2019). This study successfully met the goal of developing an integrated approach for rapid *iWUE* phenotyping. And, it used these technical advances to provide one of the largest and most comprehensive investigations of genetic and environmental variation in leaf traits that influence WUE, i.e. *g_s_* (Hatfield and Dold, 2019; Leakey et al., 2019), *A_N_* (Hatfield and Dold, 2019; Leakey et al., 2019), SD (Bertolino et al., 2019), and SLA (Zhang et al., 2009; Zhang et al., 2015). A novel element of that investigation was integration of GWAS and TWAS to identify candidate genes that can be further studied to understand and improve *iWUE* in sorghum and other C_4_ crops.

### Rapid Phenotyping

Traditional assessments of traits relating to leaf gas exchange and stomatal patterning are time and labor intensive. For example, measuring light saturated gas exchange of individual leaves can take >30 minutes (Ortiz et al., 2017; Qu et al., 2017) and manually peeling leaf epidermal samples and counting stomata via light microscopy is slow (Yates et al., 2018). Consequently, these traditional approaches are not readily amenable to large-scale assessments of genetic variation. A high-throughput phenotyping pipeline (Figure S13) was developed by integrating: (a) a rapid method of measuring leaf-level gas exchange (Figure S14; Choquette et al., 2019); b) rapid scanning of abaxial leaf surfaces and automated stomatal counting (Figures 1, S3); and c) sampling for SLA. Over 200 leaves were processed per day, facilitating phenotyping of 869 accessions replicated across two field trials in each year. This was a substantial gain in scale over previous experiments looking at similar traits in isolation (Taylor et al., 2016; Ortiz et al., 2017; Herritt et al., 2018; Lü et al., 2018; Yates et al., 2018). Our automated approach for determining SD was validated by comparisons to ground truth data (Figure 1C). And, computer-based measurement of SD in 33,355 FOV was approximately 80 times faster than counting of stomatal complexes by humans. Importantly, the efficacy of the method across a wide range of genetic and environmental variation in epidermal leaf anatomy was highlighted by the moderate-to-high heritability of SD (Figure 5). These heritability estimates were similar or higher than those previously reported (e.g. Delgado et al., 2011; Dittberner et al., 2018). A variety of machine learning methods have been developed that can identify stomata in images (e.g. Aono et al., 2019; Fetter et al., 2019; Li et al., 2019), but demonstrations of their applicability to large-scale genetic studies of the measured trait are rare (Dittberner et al., 2018) to non-existent depending on the species. Overall, this work along with Xie et al., (in review) and Bheemanahalli et al., (in review) demonstrates the utility of optical tomography and computer-vision as tools that can meet the potential for accelerating biological discovery in cereal crops.

### Genetic variation in *iWUE*, *A_N_* and *g_s_*

We detected a positive association between *A_N_* and *g_s_* (Figure 4). This is consistent with previous studies of diverse germplasm in C_4_ crops, such as maize (Choquette et al., 2019), sugarcane (Inman-Bamber et al., 2016), and switchgrass (Taylor et al., 2016). And, it affirms that accessions with greater *g_s_* achieve greater rates of *A_N_*, despite sorghum having a biochemical pump concentrating CO_2_ around Rubisco in the bundle sheath cells. However, the non-linear nature of the relationship also indicates diminishing returns from greater *g_s_* in terms of *A_N_*, leading to lower *iWUE* among accessions with the greatest *g_s_*. Selection for greater productivity in other crops has been associated with greater *g_s_* and water use (Roche 2015, Koester et al., 2016). Repeating the same strategy would not be desirable in sorghum, assuming that high productivity under water-limited conditions is a priority. Notably, the genetic variation observed in *iWUE* was more a factor of variation in *g_s_* (Figure 4C) than variation in *A_N_* (Figure 4B). Taken together these results demonstrate that enhanced *iWUE* is achieved either through low *g_s_* or through coupling high *A_N_* with moderate *g_s_*. While in the past it was suggested that WUE across C_4_ species was almost invariant (DeLucia et al., 2019), this study builds on work in sugarcane (Inman-Bamber et al., 2016) to suggest that meaningful variation does exist. Our generalized estimates of heritability for *A_N_* and *g_s_* (Figure 5) were similar to those estimated in a recent survey of the same traits in a smaller panel of grain sorghum accessions (Ortiz et al., 2017) and sufficiently high to justify targeting them as traits for selection. But, efforts to improve WUE in sorghum via direct selection on *iWUE* may inadvertently limit *A_N_* in the same way as previously observed in C3 crops (Condon et al., 2004; Leakey et al., 2019). So, understanding a broader set of component traits that influence *iWUE* will be valuable.

Contrary to theoretical expectations, and prior observations in grass crops (Miskin et al., 1972; Muchow and Sinclair, 1989; Panda et al., 2018), there was no significant correlation between SD and leaf gas exchange traits across the diverse panel of sorghum accessions (Figure 3). This was also the case in maize RILs (Xie et al., in review). But, detecting an association between SD and *g_s_* may be complicated by a strong trade-off between SD and stomatal size (Xie et al., in review). The width and length of stomatal complexes were significantly correlated with leaf gas exchange traits in maize (Xie et al., in review). Stomatal length has been observed to positively correlate with *g_s_* across rice accessions, where SD did not (Ohsumi et al., 2007). And, grain sorghum accessions selected for high or low SD alleles at a single locus display corresponding variations in *g_s_* (Bheemanahalli et al., in review). Transgenic approaches to reducing SD have reduced *g_s_* and increased *iWUE* in a number of crops (Wang et al., 2016; Hughes et al., 2017; Dunn et al., 2019; Mohammed et al., 2019). Therefore, advancing understanding of genes and traits associations underpinning SD and iWUE does have the potential to aid crop improvement efforts.

### Genetic and environmental variation in SD and SLA

SD was significantly lower in the dry growing season of 2017 than the wet growing season of 2016 (Figure 2A). The morphology and patterning of stomata can be modified in developing leaves in response to environmental cues (Lake et al., 2001; Casson and Hetherington, 2010). Lower SD would tend to limit water loss via transpiration under dry conditions, consistent with many other mechanisms that operate to sense soil water content and conserve water (Franks and Farquhar, 2001; Chaves et al., 2009). However, the response of SD to limiting water supply varies significantly among studies, species and with the intensity of drought stress (Quarrie and Jones, 1977; Hamanishi et al., 2012; Sakurai et al., 1986; Chaves et al., 2009). So, the consistent direction of response towards lower SD under drought conditions experienced in the field by this diverse sorghum population is noteworthy.

SLA was significantly greater, on average, in the drier growing season (Figure 2B), indicating an overall reduction in leaf thickness or density. SLA is well documented to demonstrate remarkable phenotypic plasticity in response to environmental stimuli (Hulshof et al., 2013; Wellstein et al., 2017). Depending on the prevailing conditions, SLA can be coupled to important functional traits, such as photosynthesis and growth rate (Pengelly et al., 2010; Liu et al., 2016; Wellstein et al., 2017; Gonzalez-Paleo and Ravetta, 2018), as well as water-use strategies and WUE (Wang et al., 2013; Scartazza et al., 2016).

Total canopy leaf area is likely to be more than sufficient to maximize light interception over most of the growing season in biomass sorghum. But, greater SLA allows greater leaf area to develop for a given investment in carbon resources. Therefore, it is possible that the greater SLA under drought conditions facilitated greater investment in other carbon sinks, such as root growth (Wellstein et al., 2017), thereby improving water uptake (Tardieu et al., 2017). Concurrently, the increase in SLA will have likely limited leaf-level carbon fixation (Xu and Zhou, 2008; Gonzalez-Paleo and Ravetta, 2018), due to a reduction in the thickness of the chloroplast-rich palisade mesophyll (Gonzalez-Paleo and Ravetta, 2018; Gotoh et al., 2018). This is reflected in the significant negative association observed between *A_N_* and SLA in 2017 (Figure 3). Despite this reduction in leaf-level *A_N_*, the side-effect of increased light penetration into the canopy may ameliorate losses in carbon gain at the canopy level (Evans and Poorter, 2001; Liu et al., 2016).

Plant height and biomass were positively correlated with the percentage change in trait values between growing seasons for both SD and SLA (Figure 3; Table S2). While many factors could contribute to this relationship, the most parsimonious explanation would be that more productive accessions generally have the greatest demand for water, exhaust available resources to the greatest extent, and then demonstrate the greatest plasticity in anatomy and physiology required to avoid further drought stress. Further work is needed to understand the adaptive value of the observed plasticity for maintenance of productivity when water is limiting. It will also be important to learn if transgenic approaches to increasing *iWUE* via lower SD constrain plasticity under drought stress.

At the genetic level, understanding of the mechanisms determining SD and SLA have not been integrated. But, accessions displaying the greatest plasticity in SD tended to be more plastic in terms of SLA as well (Figure 3). A causal link between the two traits was not explicitly tested in the current study. But, the observed correlations are consistent with previous reports of greater leaf thickness enhancing the capacity for reductions in SD under drought (Galmés et al., 2007; Xu and Zhou, 2008). Lower SLA is widely associated with greater photosynthetic capacity (Wright et al 2004). And, theory dictates that greater maximum *g_s_*, via greater SD or stomatal size, along with other aspects of hydraulic capacity in leaves should support greater exchange of water vapor for CO_2_ to be assimilated through photosynthesis (Dow et al., 2014; Henry et al., 2019). This emphasizes the need to better integrate understanding of relationships of epidermal patterning with the anatomy and function of the leaf as a whole. Consequently, candidate genes associated with variation in both SD and SLA may be of special interest (Table S7, S8).

### Combining GWAS and TWAS to identify candidate genes

The results of both GWAS and TWAS reinforced the prevailing understanding that *iWUE*, and associated leaf traits, are complex and polygenic (e.g. Des Marais et al., 2014; Ortiz et al., 2017; Dittenberger et al., 2018). As a consequence, and in common with many GWAS studies on a diverse range of traits (Zhu et al., 2008; Ortiz et al., 2017; Dittenberger et al., 2018; Kremling et al., 2019), many moderately significant associations were detected. This is consistent with individual alleles of small or moderate effect sizes segregating at moderate or low frequencies, respectively. In such cases, extra information is needed to avoid reporting false positive associations and boost confidence in the identification of candidate genes. This study provides a demonstration of the concept tested by Kremling et al., (2019) where GWAS and TWAS are combined to achieve this goal. A total of 394 unique candidate genes were identified for the set of 10 leaf traits studied. To be included in this list a gene had to be identified for a given trait in multiple independent tests for either: (1) associations of trait variation with both RNA and DNA, or (2) associations of trait variation with transcript abundance in both tissue sample types (Figure 6, Figures S4-S12, Tables S7 and S8). This was the case for 37 – 59 genes per trait, with 80 genes meeting these criteria simultaneously for 2 or 3 traits. Detailed examination of the results on a trait-by-trait basis revealed the greatest consistency in results coming from the use of Fisher’s combined test to integrate information from the GWAS with TWAS. But, there were examples where the same gene was identified from TWAS performed separately on transcriptome data from both tissue sample types (growing tip versus developing third leaf). In addition, confidence in the identification of other genes was greater because they were independently identified in both growing seasons (72 genes for SD and 69 genes for SLA) or they were identified for multiple traits resulting from independent measurements. The consistency across results for different types of traits in 2017 (213 genes for SD plus SLA; 280 genes for SD plus gas exchange traits; 288 genes for SLA plus gas exchange traits; Figure 7B) was than higher than for across growing seasons. But, this does not seem surprising given the difference in water availability between the two years and the potential for genotype x environment interactions. Confirmation of a role for these genes in driving variation in *iWUE* and related traits will still require a reverse genetics approach performed on a gene-by-gene basis. But, a significant number of the candidate genes identified are orthologs of genes in *A. thaliana* that have functions linked in some way to iWUE, *A_N_*, *g_s_* or leaf development and anatomy (Table S9). And, the preponderance of genes with associations between trait variation and transcript abundance may indicate that regulatory variation is a more common driver of genetic variation than sequence variants. It is worth noting that TWAS was performed using transcriptome data generated from plants in controlled conditions, as opposed to the field where phenotyping for GWAS was performed. Despite this, the molecular control of the physiological processes of interest is well conserved, which is reflected in the overlap of genes identified by association to RNA and DNA.

### Candidate genes underlying variation in gas exchange and SD

In terms of genes identified underlying variation in *g_s_*, but not *A_N_*, *MAPK4*, identified via GWAS for *g_s_*, represents a particularly promising candidate (Table S9). *MAPK4* is a well characterized disease resistance protein (Berriri et al., 2012), but evidence from aspen (Witoń et al., 2016) and agave (Sara et al., 2020) demonstrate a role for *MAPK4* in the regulation of *g_s_* and stomatal development, which is in line with the identification of *MAPK4* via TWAS for SD in GP tissue also.

We did not observe strong associations between SD and gas exchange traits. This is possibly due to the restricted variation in SD in 2017. However, it is also likely due to the importance of further uncharacterized stomatal and mesophyll components for regulating gas exchange (Bertolino et al., 2019; Lawson and Matthews, 2020). Despite this, it is well understood that significantly manipulating SD can improve WUE in many species (Bertolino et al., 2019). Consequently, the candidate genes underlying SD identified in this study represent a starting point for future crop improvement to this end. Moreover, elucidating these candidates for their role in regulating SD may form the foundations of understanding stomatal development pathways in C_4_ grasses.

Candidate SD genes with known roles in stomatal development included *MUTE*, which was identified via the Fisher test employing GP transcriptome data (Figure 6, Table S9). *MUTE* is a bHLH transcription factor that acts as a stoma-fate master regulator in dicots and monocots (Pillitteri et al., 2008; Raissig et al., 2017).

Further genes with demonstrated roles in stomatal development identified within this set included *KCS1* which controls stomatal patterning relative to CO_2_ concentration (Gray et al., 2000) and *HB-7* which regulates stomatal size relative to water availability (Ré et al., 2014) (Figure 6, Table S9). Putative SD candidates in this gene set included a cell wall expansion-type protein (*EXPB2* (Marowa et al., 2016), an ABA-sensitive MAP KINASE (*RAF10* (Lee et al., 2015)), the *PAP10* purple acid phosphatase (Hepworth et al., 2016), and an asparagine-rich protein (*NRP*) that is documented to positively regulate the expression of *CRY2* (Zhou et al., 2017), a blue light receptor which in turn increases stomatal index (Kang et al., 2009) (Figure 6, Table S9). Further genes with known roles in stomatal development identified via alternative mapping approaches included *AMP1* (Shi et al., 2013; López-García et al., 2020), *ATE1* (Movahedi, 2013; Vicente et al., 2019), and *TED5* (Tossi et al., 2014; Zoulias et al., 2020). Additionally, through mapping for SD we identified multiple genes with putative and known roles in stomatal behavior, e.g. the *CLC-C* anion transporter (Jossier et al., 2010), and ABA responsiveness, e.g. *FRS5* (Ma and Li, 2018), that represent interesting targets for further study.

*CRR23* was identified via all mapping approaches for *A_N_* (Table S9) and is known to be critical for stabilizing the chloroplast NAD(P)H dehydrogenase complex, thereby facilitating photosynthetic electron transport (Shimizu et al., 2008). The importance of this complex in controlling the observed variation in *A_N_* was further highlighted by the identification of the *AOX1a* gene via multiple *A_N_* mapping approaches (Table S9). *AOX1a* is well demonstrated to play a key role in electron transport and balancing the redox state of cellular NAD(P)H pools, thereby facilitating efficient photosynthetic functioning (Vishwakarma et al., 2014; Podgórska et al., 2020). Combined GWAS and TWAS for *A_N_* also identified genes with demonstrated roles in chloroplast biosynthesis. For example, *PDS3* (Table S9) is a key component of retrograde signaling during chloroplast development. Indeed, *pds3*-mutants display an albino phenotype (Foudree et al., 2010). *PDS3* was also identified via mapping for *g_s_* and SD, which is interesting since *PDS* genes have been implicated in ABA biosynthesis and the control of stomatal opening (Chao et al., 2014). *PIF3* represents a further candidate in this vein. *PIF3 w*as identified via mapping for *A_N_* and *g_s_* (Table S9). *PIF3* is a light-dependent transcriptional repressor of genes involved in chlorophyll biosynthesis and further photosynthetic processes (Liu et al., 2013). Additionally, the closely related *PIF4* gene has been demonstrated to regulate the expression of *SPEECHLESS* (*SPCH*), a master regulator of stomatal development (Casson et al., 2009; Lau et al., 2018), thereby hinting at a possible role in regulating *g_s_*. Additionally, mapping for *A_N_* identified *PIP1;4* (Table S9), which is an aquaporin that regulates the permeability of the plasma membrane to CO_2_, thereby mediating CO_2_ transport for photosynthesis (Li et al., 2015).

## CONCLUSION

This study demonstrates the application of novel high-throughput phenotyping tools with combined GWAS/TWAS to study the genetic basis for a challenging set of complex traits related to *iWUE* in a model C_4_ crop. In doing so, it revealed heritable variation in multiple traits that selection could act upon to improve performance under water limited conditions. In addition, it highlights the central role that SLA may play as an allometric trait that is associated with broad genetic and environmental variation in SD, leaf photosynthetic gas exchange and plant productivity. Lastly, genomic and transcriptomic variation across this diversity set were leveraged to identify multiple candidate genes with known and putative roles for key WUE traits.

## MATERIALS AND METHODS

### Germplasm and experimental design

869 previously described biomass sorghum accessions (Valluru et al., 2019; Dos Santos et al., 2020) were used in this study (Figure S1; Table S1). All lines were grown during 2016 and 2017 across two field sites in Central Illinois (Savoy, IL), where experiments were sown in late May and harvested in late October. Lines were grown according to an augmented block design as reported previously (Valluru et al., 2019; Dos Santos et al., 2020).

### High throughput phenotyping pipeline for WUE-associated traits

The youngest fully expanded leaf of two plants randomly selected from the middle two rows of each plot were excised slightly above the ligule between September 5 and 14, 2016. Damaged leaves were avoided. Excised leaves were immediately placed in a bucket, with the cut surface submerged under water. In the laboratory, three 1.6cm leaf discs were collected from each leaf while avoiding the midrib. Leaf discs were immediately transferred to an oven set at 60°C for two weeks. The dry mass of leaf discs was determined and specific leaf area (SLA) was calculated as the ratio of fresh leaf area to dry leaf mass (cm^2^ g^−1^). The SLA data collected in 2016 were previously reported (Valluru et al., 2019).

A leaf tissue strip approximately 1cm x 3cm in area was also cut from the adjacent portion of the leaf from where the leaf discs were collected. Leaf strips were marked to distinguish the abaxial side, inserted into 2 mL screw cap tubes and flash frozen in liquid nitrogen and stored at −80°C. ~150 leaf strip samples were moved to −20°C during active microscopy. Leaf samples were removed from the −20℃ freezer and affixed to a microscope slide using double-sided tape with the abaxial side facing up. The surface topography of leaf surfaces was measured using two Nanfocus μsurf explorer optical topometers (Nanofocus, Oberhausen, Germany) at 20x magnification with a standardized area of 800 × 800 μm. The upper and lower z-scale limits being set manually for each FOV to ensure all stomata were in focus. The abaxial surface topography was measured at 4-6 randomly selected points producing 4-6 fields of view (FOV).. Measurements were saved in the .nms file format and automatically transferred for automated stomatal counting via a CyVerse-enabled data pipeline (Goff *et al.* 2011).

In 2017, leaf-level gas exchange of all accessions was measured in addition to sampling for SLA and tissue for stomatal imaging. The field was divided into four quartiles based on height measured in previous growing seasons. Each quartile was sampled over a 4- or 5-day period. On each measurement day, 200-232 leaves (2 leaves from 100-116 plots) were harvested pre-dawn as described for the 2016 SLA and SD leaf sampling. Upon returning the leaves to the lab, stable rates of light-saturated gas exchange were measured by following the experimental protocol described previously for maize (Choquette *et al.* 2019). Stable rates of net photosynthetic assimilation of CO2 (*A_N_*), stomatal conductance (*g_s_*), intrinsic water use efficiency (iWUE), and the ratio of intracellular and atmospheric CO2 (*c_i_*/*c_a_*) were obtained by averaging data from the last two minutes of a four-minute autolog program (Figure S14). After the measurements of leaf-level gas exchange, the area of the leaf contained within the cuvette was marked and used for sampling for leaf discs and tissues strips for subsequent measurements of SLA and stomatal imaging as described above for the 2016 sampling campaign. A flow chart describing this pipeline is provided in Figure S13.

### Automated stomatal counts

Image processing and machine learning methods were combined to produce a software tool that automatically detected stomata in 33,355 grayscale images of sorghum leaf surfaces. Constructing the method required a set of training data based on circular image disks 80 pixels in diameter centered where human experts had registered the locations of stomata in many 512×512 raw images. Each 80-pixel disk was subjected to circular fast Fourier transformation (FFT) to produce a radial series of phase and amplitude values that proved to be predictive of stomata. The radial FFT results were recast by principal components analysis (PCA) into a lower dimensional form that served as the feature set used to train nine different machine learning methods. The nine methods were: an Artificial Neural Network, Linear Discriminant Analysis, a Convolution Neural Network, three Generalized Linear Model‘s two Regularization (Ridge and Lasso) and one without, Partial Least Squares Regression, Stepwise Linear Regression and a Decision Tree. Each method produced a version of the original image in which each pixel value was a probability of that location belonging to a stoma. Next, a fusion process filtered and combined the independent probability maps such that local probability peaks in excess of a height threshold optimally coincided with the locations of human-verified stomata. Specifically, a Nelder-Mead optimization process adjusted the filter and threshold parameters to maximize the agreement between the machine-labeled stomata and the human-identified stomata as quantified by the Matthews correlation coefficient. Figure S3 shows an overview of the method. The analyses were implemented in the Matlab programming environment and deployed on a high throughput computing resource with jobs scheduled by HTCondor (Thain, Tannenbaum & Livny 2005). The machine learning and optimization processes (i.e. layers) were subsequently trained and tuned accordingly.

Each FOV from a stomatal imaging sample produced a stomatal count value. The stomatal count values were divided by the area of the images (0.64mm^2^) to give stomatal density (SD). The median of the 4-6 SD estimates were calculated for each sample and used for subsequent analyses. To benchmark the efficiency of the automated stomatal counting, we manually counted and estimated stomatal density for 227 randomly selected samples, which represented 1056 individual FOV. A linear model predicting manually counted stomatal density from automatic stomatal density was subsequently fit.

### Plant height and biomass measurements

In 2016 and 2017, a single representative plant in each plot was measured for plant height as described and reported previously (Valluru et al., 2019; Dos Santos et al., 2020). For this study, plant height on 105 days after planting (DAP) was used for comparative analyses since it showed the greatest heritability of all days measured. In 2017, plants were harvested, and above ground biomass was measured and calculated as dry t ha-1 as previously described and reported (Dos Santos *et al.* 2020)

### Statistical models and heritability

For each trait, we fitted a linear mixed model using the ASReml-R v4.0 package (Butler *et al*., 2018). The appropriate model was chosen based on the Akaike information criterion (AIC) and the diagnostic plots. As different covariables were evaluated along with each phenotype, the final model varied in each case (Table S10). The general model used was as follows:

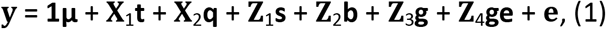

where **y**(n × 1) is the vector of phenotypes for j environments (year × location combination) with 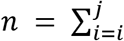 *n_i_*; **1**(n × 1) is a vector of ones; **μ** is the overall mean; **X**_1_ (n × j) is the incidence matrix associated with the vector of fixed effect environments **t**(j × 1); **X**_2_ (n × v) is the incidence matrix associated with the vector of fixed effect covariates **q** (v × 1) (see supplementary material for details on the number of fixed effects covariates used in each model, if any); **Z**_1_ (n × f) is the incidence matrix associated with the vector of random effect set within environment **s** (f × 1) with *s* ~ *MVN*(0, *I_f_* ⊗ ***S***); **Z**_2_ (n × r) is the incidence matrix associated with the vector of random block within set within environment effects **b**(r × 1) with *b* ~ *MVN*(0, *I_r_* ⊗ *B*); **Z**_3_ (n × l) is the incidence matrix associated with the vector of random genotype effects **g**(l × 1) with *g* ~ *MVN*(0, *I*_l_ ⊗ *G*); **Z**_4_ (n × w) is the incidence matrix associated with the vector of random genotype-by-environment effects **ge** (w × 1) with *ge* ~ *MVN*(0, *I_w_* ⊗ *K*); and **e** (n × 1) is the vector of residuals with 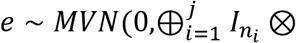 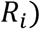. The matrices **S**, **B**, **G**, **K**, and **R** are the variance–covariance matrices for set within environment, block within set within environment, genotype, genotype-by-environment, and residual effects, respectively. For each genotype, we obtained predictions from model 1 and these were used for downstream analysis. The generalized heritability was estimated as proposed by (Cullis et al., 2006).

### RNA-seq analysis

A subset of the full diversity panel was grown under controlled experimental conditions for 3’ RNAseq analysis of genes potentially involved regulating leaf development, including stomatal patterning. The abundances of transcripts for orthologs of known stomatal patterning genes were initially screened across 3-7 separate tissues at each of 4 developmental stages during the day and night in six accessions. On the basis of that screen, the base of leaf three and the shoot growing point at the 3-leaf stage were targeted for sample collection during the day from a subset of 229 accessions from the full population. Samples were processed and expression data was generated from libraries using a pipeline and parameters similar to Kremling et al. (2019). Briefly, reads were trimmed using Trimmomatic (version 0.32) to remove adapter sequences in relation to *Illumina chemistry* and sequencing errors. Next, trimmed reads were aligned to the sorghum reference genome (version 3.1.1) using the splice-aware aligner, STAR (Spliced Transcripts Alignment to a Reference) (version 2.4.2). Feature counts were then generated using HTSeq (version 0.6.1) from previously generated alignment files. Finally, count normalization was performed using the R package, DESeq2 via size factor estimation.

### GWAS

For conducting GWAS, we imputed the 100,435 GBS SNPs from (Dos Santos et al., 2020) using as reference panel the whole-genome re-sequencing dataset of 5,512,653 SNPs published by Valluru et al. (2019). The untyped genotypes were imputed and phased into haplotypes using Beagle 4.1 using a default window size of 50,000 SNPs and an Ne = 150,000. After the imputation, SNPs with allelic R squared < 0.5 and minor allele count below 20 where removed, resulting in a total of 2,327,896 SNPs. Additionally, we pruned SNPs in high linkage disequilibrium (r^2^> 0.9) using Plink options “--indep-pairwise 50 10 0.9”. The final data set consisted of 454,393 SNPs scored in 836 sorghum lines.

The association analysis was conducted using the unified MLM (Yu et al. 2006) implemented in the software GEMMA (Zhou and Stephens, 2012). For that, predicted values obtained from model 1 were normal quantile transformed as done in (Zhou and Stephens, 2014). We used the Bayesian information criteria (BIC; Schwarz, 1978) to select the appropriate number of principal components (PCs) to account for population structure. We tested models with 0-10 PCs estimated from TASSEL 5 (Bradbury et al., 2007). The best model did not include any PC. Relatedness was controlled for by a kinship matrix obtained from TASSEL 5 using the default method (Endelman and Jannink, 2012).

### TWAS and combined GWAS-TWAS

A transcriptome-wide association study (TWAS) was performed on a subset (229) of the total accessions (Valluru et al. 2019) and conducted using TASSEL (version 5.2.5). Before mapping, covariates were generated from multiple sources. 10 hidden factors were calculated using probabilistic estimation of expression residual (PEER) factors for each individual tissue (Stegle et al. 2012). Additionally, 5 genetic principal coordinates (PC) were calculated from prior genotype data (Valluru et al. 2019). Genes that were expressed in at least half of the individual lines were used within each tissue set. A general linear model was fit individually for each phenotype and every combination of expressed gene value across individuals after adjusting for PC and PEER factor covariates.

TWAS-GWAS combined p-values were calculated in a similar fashion as described by Kremling et al. (2019). Briefly, p-values of the top 10 percent significant GWAS SNPs were assigned to their nearest gene. Assigned GWAS p-values were then combined with their respective TWAS p-values via the Fisher’s combined test as using the sumlog function within the R package, metap.

To further explore the results of all genetic mapping approaches, we queried commonality between specific gene sets, e.g. genes identified for SD and SLA, SD genes common to 2016 and 2017, Genes identified via GWAS and Fisher’s combined test, etc. (Figure 7B). For these comparisons, the total number of possible shared genes between any two gene sets was determined.

### Candidate gene identification

For each GWAS result, the top 0.1% of SNPs based on −log_10_(*p-value*) were identified. The linkage disequilibrium (LD) blocks these SNPs associated with was determined and all genes within these LD blocks or spanning their borders were extracted. LD blocks were estimated based on the method proposed by (Gabriel, 2002) and implemented in PLINK (Chang et al., 2015). For this, we used the option −-blocks, with a window of 200 kb and default values for D-prime’s confidence interval (0.7;0.98).

For the TWAS and the combined GWAS-TWAS, the top 1% of genes based on −log_10_(*p-value*) were identified for each result. A list of candidate genes with known or putative roles in associated traits was determined based on overlap between different mapping approaches and/or traits.

### Additional statistical analysis and figure generation

All further statistical analysis and figure generation was performed within the R environment (R Core Team, 2017). Change in SLA and SD across the two growing seasons was calculated at the accession level as percentage change using adjusted means. SLA increased in all but two accessions across the two years, thus percentage change in SLA was calculated as: (2017_SLA_-2016_SLA_) / 2017_SLA_ * 100. SD decreased in all accessions across the two years, thus percentage change in SD was calculated as: (2016_SD_-2017_SD_) / 2016_SD_ * 100. Tests for associations between all pairwise trait interactions were performed using adjusted means from all models and the Pearson’s product moment correlation coefficient. Pairwise interactions between specific leaf-level gas exchange traits were further investigated by fitting second-order polynomial equations between traits. Except for Manhattan plots, all figures were generated using the R package ggplot2 (Wickham, 2016). Manhattan plots for all genetic mapping visualization were generated using the R package qqman (Turner, 2017).

## Acknowledgments

We thank Victoria Scaven, Anna Dmitrieva, Dylan Allen, Aishwarya Kammala, and Eric Peterson for help with high throughput phenotyping data collection. We thank Jose Antonio Cumbra, Lauren Murphy, and Abigail Garcia for assistance with the manual stomatal counting. We thank Charles Pignon for helpful discussions and assistance with data analysis. The information, data, or work presented herein was funded in part by the Advanced Research Projects Agency-Energy (ARPA-E), U.S. Department of Energy, under Award Number DE-DE-AR0000661. The views and opinions of the authors expressed herein do not necessarily state or reflect those of the United States Government or any agency thereof.

## Supplemental Figure Legends

Supplemental figure 1. Map showing the point of origin of all accessions employed in this study. Accessions are colored according to race. The size of the points indicates the number of accessions from common origins. The inset biplot shows the genetic structure of all accessions according to principle component analyses.

Supplemental figure 2. Combined line and bar plots of key climate parameters during the growing season of (a) 2016 and (b) 2017. Each plot describes the daily precipitation (blue bars) and maximum daily temperature (red line) from the last week of May until the last week of September. For each year, the total precipitation and average maximum temperature are provided.

Supplemental figure 3. Overview of the custom stomata counting software method. (a) Raw image of sorghum leaf surface with a stoma circled. (b) Enlargement of a disc containing a stoma. A FFT process extracted image features at a series of 25 radii, only three of which are indicated with magenta circles. PCA decomposition of the FFT phase and amplitude values extracted from expert-labeled stomata discs were used to train 9 different machine learning algorithms, each of which produced a pixel value representing probability of belonging to a stoma. (c) Example of a stoma probability map from one of the machine learning methods. A fusion process combined the 9 probability maps and optimized filter parameters to produce a binary classifier that assigned each pixel to a stoma or not-stoma class. (d) A final binary stomata map from which counts were obtained.

Supplemental figure 4. Manhattan plot demonstrating results of GWAS, TWAS, and fisher combined GWAS and TWAS of stomatal density (SD; 2017 model)

Supplemental figure 5. Manhattan plot demonstrating results of GWAS, TWAS, and fisher combined GWAS and TWAS of stomatal (SD; Joint model)

Supplemental figure 6. Manhattan plot demonstrating results of GWAS, TWAS, and fisher combined GWAS and TWAS of specific leaf area (SLA; 2016 model)

Supplemental figure 7. Manhattan plot demonstrating results of GWAS, TWAS, and fisher combined GWAS and TWAS of specific leaf area (SLA; 2017 model)

Supplemental figure 8. Manhattan plot demonstrating results of GWAS, TWAS, and fisher combined GWAS and TWAS of specific leaf area (SLA; joint model)

Supplemental figure 9. Manhattan plot demonstrating results of GWAS, TWAS, and fisher combined GWAS and TWAS of stomatal conductance (*g_s_*)

Supplemental figure 10. Manhattan plot demonstrating results of GWAS, TWAS, and fisher combined GWAS and TWAS of net photosynthesis (*A_N_*)

Supplemental figure 11. Manhattan plot demonstrating results of GWAS, TWAS, and fisher combined GWAS and TWAS of intrinsic water use efficiency (*iWUE*)

Supplemental figure 12. Manhattan plot demonstrating results of GWAS, TWAS, and fisher combined GWAS and TWAS of the ratio of intracellular to atmospheric CO2 concentrations (*c_i_*/*c_a_*)

Supplemental figure 13. Flow chart demonstrating the phenotyping pipeline employed in this study during the 2017 field season. In 2016, leaf-level gas exchange data were not collected.

Supplemental figure 14. Scatter plots of representative examples of leaf-level gas exchange logged during the phenotyping pipeline. Data points for gas exchange were logged every four seconds for four minutes. The mean for the final 15 data points (highlighted in red) was calculated to achieve sample values for (a) net photosynthesis (*A_N_*), (b) stomatal conductance (*g_s_*), (c) ratio of intracellular to atmospheric concentrations of CO2 (*c_i_*/*c_a_*), and (d) intrinsic water use efficiency (*iWUE*).

Supplemental figure 15. Comparison of plant height at 120 days after planting in 2016 and 2017 (Data from: (Dos Santos et al., 2020)(a) Histograms showing variation in genotypic means of plant height in 2016 and 2017. (b) Correlation between genotypic means for plant height in 2016 and 2017.

## Supplemental Tables

Supplemental table 1. List of accessions comprising this study.

Supplemental table 2. Pearson’s correlation coefficients and *p-values* for associations between all traits

Supplemental table 3. Top 0.1% of SNPs identified through GWAS for each trait

Supplemental table 4. Genes with linkage disequilibrium of the top 0.1% SNPs identified through GWAS for each trait

Supplemental table 5. Top 1% of genes identified through TWAS for each trait and each tissue type

Supplemental table 6. Top 1% of genes identified through Fisher combined GWAS-TWAS for each trait and each tissue type

Supplemental table 7. Tall version of genes identified in the top sets of all mapping approaches. The mapping approach and trait for which they were identified is listed.

Supplemental table 8. Wide version of genes identified in the top sets of all mapping approaches. The mapping approach and trait for which they were identified is listed.

Supplemental Table 9. List of candidate genes with known or putative roles associated to the traits for which they were identified

Supplemental table 10. List of models mixed models used in this study to obtain adjusted genotypic means for all traits

Supplemental table 11. Adjusted genotypic means for all traits comprising this study

